# Sanctions, partner recognition, and variation in mutualism

**DOI:** 10.1101/049593

**Authors:** Jeremy B. Yoder, Peter Tiffin

## Abstract

Mutualistic interactions can be stabilized against invasion by non-cooperative individuals by putting such “cheaters” at a selective disadvantage. Selection against cheaters should eliminate genetic variation in partner quality — yet such variation is often found in natural populations. One explanation for this paradox is that mutualism outcomes are determined not only by responses to partner performance, but also by partner signals. Here, we build a model of coevolution in a symbiotic mutualism, in which hosts’ ability to sanction non-cooperative symbionts and recognition of symbiont signals are determined by separate loci, as are symbionts’ cooperation and expression of signals. In the model, variation persists without destabilizing the interaction, in part because coevolution of symbiont signals and host recognition is altered by the coevolution of sanctions and cooperation, and vice-versa. Individual-based simulations incorporating population structure strongly corroborate these results. The dual systems of sanctions and partner recognition converge toward conditions similar to some economic models of mutualistic symbiosis in which hosts offering the right incentives to potential symbionts can initiate symbiosis without screening for partner quality. These results predict that mutualists can maintain variation in recognition of partner signals or in the ability to sanction non-cooperators without destabilizing mutualism, and reinforce the notion that studies of mutualism should consider communication between partners as well as the exchange of benefits.

## Introduction

Mutually beneficial interactions between species pose two related conundrums. First, how are mutualisms maintained in the face of the apparent advantages to individuals who accept resources or services but provide none in return? And second, given a mechanism that prevents the evolution of non-cooperative participants, why do members of interacting species vary in mutualistic quality?

The first conundrum may be solved through selective dynamics that offset cheaters’ advantage. Mutualists might avoid or discontinue interaction with cheaters (Trivers 1971; Axelrod and Hamilton 1981; Foster et al. 2006), or might reduce or cut off rewards provided to them (Bull and Rice 1991; West et al. 2002a; West et al. 2002b; Sachs et al. 2004; Akçay and Simms 2011). Some authors have differentiated the former approach as “partner choice,” though both are in some sense *sanctions* that actively deny the full benefit of mutualism to cheaters and avoid the full cost of interacting with them. By contrast, in *partner fidelity feedback*, cooperative partners receive greater rewards without any active “decision” by the reward-providing species, simply because healthy individuals produce more rewards (Doebeli and Knowlton 1998; Weyl et al. 2010; Frederickson 2013) or because rewards are only accessible to cooperative partners (Archetti et al. 2011a; Archetti et al. 2011b). Each of these mechanisms ensure that non-cooperators are at a long-term fitness disadvantage even if they have an advantage over cooperators in the short term.

Such anti-cheating mechanisms have been found in many mutualisms. Legumes can cut off resources to root nodules in which rhizobial bacteria do not fix nitrogen (Kiers et al. 2003; Batstone et al. 2017) and can scale these sanctions quantitatively to reduce support for less-productive nodules (Kiers et al. 2006). In the obligate brood pollination mutualisms of yuccas and figs, in which pollinators lay eggs in flowers in the course of delivering pollen, host plants abort flowers that are too badly damaged by pollinator oviposition or that receive poor-quality pollination (Pellmyr and Huth 1994; Jandér and Herre 2010). Ant-protected shrubs reduce the growth of domatia, structures that house protective ants, on branches that suffer herbivore damage, and this may be a sanctioning response to poor protection by the ants (Edwards et al. 2006a), or may be an example of partner fidelity feedback (Weyl et al. 2010).

These solutions to the first conundrum of mutualism create the second conundrum. Partner choice, sanctions, and partner fidelity feedback should all lead to fixation of cooperative genotypes (Axelrod and Hamilton 1981; Doebeli and Knowlton 1998; West et al. 2002a; West et al. 2002b). Similarly, if interacting mutualists maximize their own fitness by matching each other, coevolution should reduce diversity in both interacting species (Kiester et al. 1984; Kopp and Gavrilets 2006; Yoder and Nuismer 2010). Nevertheless, genetic variation in partner quality is widely observed in natural populations of mutualists (Heath and Stinchcombe 2014; Jones et al. 2015), including in many interactions where mutualism-stabilizing mechanisms have been studied directly, such as rhizobial bacteria and their legume hosts (Simms and Taylor 2002; Heath and Tiffin 2009), mycorrhizal fungi (Hoeksema 2010), ant bodyguards (Ness et al. 2006), and brood pollinators (Pellmyr and Huth 1994; Herre and West 1997; Holland et al. 1999).

Coevolutionary dynamics that can maintain genetic variation in interacting species are known — not from mutualism, but from antagonistic interactions. Host-parasite interactions or competition can either create frequency dependent selection on interacting species (e.g., Dieckmann et al. 1995; Agrawal and Lively 2002), or select for one species to be less well matched to the other (Sasaki 2000; Nuismer and Otto 2005; Kopp and Gavrilets 2006; Yoder and Nuismer 2010). These dynamics are well documented in biological systems in which host defensive responses are activated by recognition of molecules expressed by parasites or pathogens (reviewed by Dybdahl et al. 2014; Nuismer and Dybdahl 2016).

Mutualistic interactions also require processes of recognition and signal exchange. Many brood pollinators respond to complex, host-species-specific floral scents that are not directly related to rewards offered by hosts (Svensson et al. 2005; Okamoto et al. 2007; Soler et al. 2011; Svensson et al. 2016). Host plant volatiles also guide the colonizing queens of plant-protecting ant species and direct the activity of ants’ patrols (Edwards et al. 2006a; Edwards et al. 2007; Schatz et al. 2009). Legumes recognize and respond to signals and identifying surface proteins expressed by rhizobia and mycorrhizal fungi (Oldroyd et al. 2011). Immune recognition responses help determine the assembly of animals’ microbiomes (Pflughoeft and Versalovic 2012; Cullender et al. 2013; Mutlu et al. 2014). Such signaling and recognition factors in interacting mutualists may coevolve in very different ways from traits governing mutualistic performance, and coevolution of signals and responses to them may affect the coevolution of benefits exchanged.

We hypothesize that coevolving partner communication maintains variation in mutualism outcomes even as sanctions prevent the breakdown of mutualism. Here, we test this hypothesis with models of a mutualism in which outcomes are determined by (1) sanctions against non-cooperative individuals, (2) recognition of signals that are separate from symbiont quality, or (3) sanctions against non-cooperators paired with recognition of signals. We first present analytic models of allele frequency dynamics within a population of two interacting species, then use individual-based coevolutionary simulations to examine a wider range of parameters, and to examine how coevolution in the different models shapes geographic variation as well as local diversity. We find that sanctions alone maintain mutualism without variation, while recognition alone maintains variation but not the mutualism. Incorporating both sanctions and recognition can maintain the mutualism as well as variation in the outcomes of mutualists’ interactions.

## Methods

We model a mutualism with outcomes determined by sanctions against non-cooperative partners, by recognition of partner signals separate from cooperation, or by both sanctions and recognition. The model is inspired by symbiotic mutualisms such as brood pollination (Pellmyr and Huth 1994; Jandér and Herre 2010) or nutrient symbioses like the legume-rhizobium mutualism (Kiers et al. 2003), in which partly or fully free-living individuals of one species provide a benefit to members of another species, which provide rewards in return. As in many biological systems the first of these species, the symbiont, is the one considered a potential “cheater,” while the second species, the host, exerts sanctions against such individuals. For the purposes of this model, we consider that “sanctions” refer to any response by hosts such that they pay a reduced cost of hosting a symbiont that provides no benefit, and deny the full benefit of symbiosis to that non-cooperator. Many authors have used “partner choice” to refer to mutualists ceasing interaction with non-cooperative individuals (e.g., Trivers 1971; Axelrod and Hamilton 1981; Foster et al. 2006); the sanctions in our model can encompass such partner choice (following, e.g., Kiers et al. 2003; Segraves 2003; Jander and Herre 2016).

In all three models, we assume that hosts and symbionts encounter each other at random and interact, and that each species *i* receives a benefit *B*_*i*_ and pays a cost *C*_*i*_ of interaction. For both host and symbiont we assume that fitness is equal to 1 + *P*_*jk*_, where *P*_*jk*_ is the payout (i.e., net benefit) from the interaction of an individual with genotype *j* interacting with a member of the other species with genotype *k*. Payout is determined by host and symbiont genotype and by the possible benefit (*B*_*i*_) and the cost (*C*_*i*_) of interaction. We assume that *B*_*i*_ *> C*_*i*_, which restricts our analysis to conditions under which both partners can potentially receive a positive payout from the interaction.

### Analytic models

We first derive three analytic models of coevolving hosts and symbionts, which track allele frequencies in both species within a single population. Full details of model derivations and evaluation are available as Mathematica notebooks, deposited in the Dryad Digital Repository:http://dx.doi.org/10.5061/dryad.p2s02 (Yoder and Tiffin 2017).

#### Host sanctions

First, consider a model of host sanctions against non-cooperative symbionts, in which symbiont cooperation and hosts’ ability to sanction are each determined by a single biallelic locus. Symbionts with the *M* allele at a *cooperation* locus cooperate; symbionts with the *m* allele do not. Cooperative symbionts pay a cost of symbiosis, *C*_*S*_, and receive a benefit, *B*_*S*_, while non-cooperative symbionts receive the benefit but pay no cost. A host interacting with a cooperative symbiont pays a cost of hosting symbionts, *C*_*H*_, and receives a benefit of symbiosis, *B*_*H*_. Hosts interacting with non-cooperative symbionts receive no benefit, but pay the cost unless they are able to sanction.

Hosts with the *H* allele at a *sanctions* locus are able to stop interaction with a non-cooperating symbiont with effectiveness *ω*; hosts with the *h* allele are not able to do so. The term *ω* determines the degree to which sanctioning hosts are able to avoid paying the costs of hosting non-cooperating symbionts and deny them the benefit of symbiosis. If *ω* = 1, sanctioning hosts suffer no cost of hosting non-cooperators and the non-cooperators receive no benefit; if *ω* = 0, sanctions have no effect, so that hosts pay the full cost of symbiosis and non-cooperators receive the full benefit (Axelrod and Hamilton 1981; Ohtsuki 2010). We do not include a separate term for a cost paid by hosts when they apply sanctions, but a cost is implicit if sanctions are less than fully effective (*ω <* 1) and there is a non-zero cost of hosting symbionts (*C*_*H*_ *>* 0). This parallels empirical systems in which sanctions cut off interaction after an initial investment, such as legumes that initiate nodulation with low-quality rhizobia only to reduce investment in less-productive nodules (Kiers et al. 2006), or yuccas and figs that invest in flowers, but abort them if they are too badly damaged by seed-feeding pollinators (Pellmyr and Huth 1994; Jandér and Herre 2010).

As noted above, host and symbiont fitness are equal to 1 + *P*_*jk*_, where *P*_*jk*_ is the payout from an individual with genotype *j* interacting with a member of the other species with genotype *k*, as determined by the possible benefit (*B*_*i*_) and the cost (*C*_*i*_) of interaction to each species *i*, and by the effectiveness of sanctions *ω* (Table 1).

**Table 1:** Host and symbiont payouts under the model of host sanctions.

| Symbiont | Host |  |
| --- | --- | --- |
|  | H | h |
| *Host payout* |  |  |
| M | $B_H - C_H$ | $B_H - C_H$ |
| m | $-(1 - \omega)C_H$ | $-C_H$ |
| *Symbiont payout* |  |  |
| M | $B_S - C_S$ | $B_S - C_S$ |
| m | $(1 - \omega)B_S$ | $B_S$ |

We can then derive the per-generation change in the frequency of the host’s *H* allele:

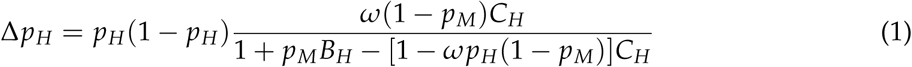

And the symbiont’s *M* allele:

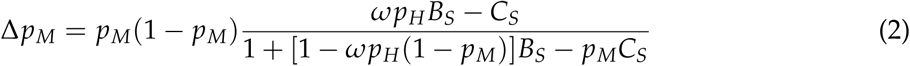

#### Partner recognition

Next, consider a model of partner recognition, in which hosts only interact with symbionts expressing a signal compatible with the hosts’s recognition genotype, and symbiont signals are determined by a locus independent of the locus that determines whether symbionts cooperate. This is essentially a “matching alleles” infection genetics model of the type used to study host-parasite interactions (Agrawal and Lively 2002; Nuismer 2017).

As in the sanctions model, symbionts cooperate if they have the *M* allele at the cooperation locus, and do not if they have the *m* allele. Symbionts also carry either a *S* allele or an *s* allele at a *signaling* locus. Hosts have no ability to sanction non-cooperating symbionts; instead they carry either a *R* or *r* allele at a *recognition* locus. Hosts with the *R* allele initiate symbiosis only with symbionts carrying the *S* signaling allele, hosts with the *r* allele initiate symbiosis only with symbionts carrying *s*. With no ability to sanction, hosts’ payouts are determined solely by whether symbionts carrying compatible signaling alleles are also cooperative (Table 2).

**Table 2:** Host and symbiont payouts under the model of partner recognition.

| Symbiont | Host |  |
| --- | --- | --- |
|  | R | r |
| *Host payout* |  |  |
| MS | $B_H - C_H$ | 0 |
| Ms | 0 | $B_H - C_H$ |
| mS | $-C_H$ | 0 |
| ms | 0 | $-C_H$ |
| *Symbiont payout* |  |  |
| MS | $B_S - C_S$ | 0 |
| Ms | 0 | $B_S - C_S$ |
| mS | $B_S$ | 0 |
| ms | 0 | $B_S$ |

An exact analytic examination of equilibria in this model is impractical. However, if we assume that the costs and benefits of the interaction are small (Nuismer et al. 2010; Yoder and Nuismer 2010), that the effects of the symbiont cooperation (M) and signaling (S) loci are therefore not strongly epistatic, and that there is free recombination between symbiont loci, then alleles at these loci should remain in quasi-linkage equilibrium (QLE) conditions (Barton and Turelli 1991; Kirkpatrick et al. 2002). With these assumptions, we can approximate change in the frequency of the host *R* allele as

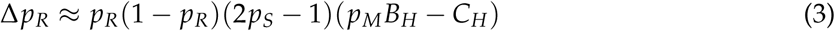

the symbiont’s *M* allele as

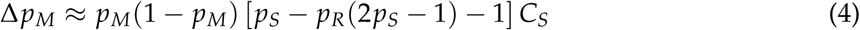

and the symbiont’s *S* allele as

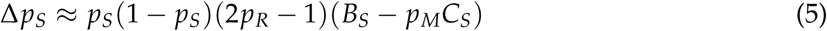

We also calculate an approximate per-generation rate of change in LD between alleles at the two symbiont loci. However, under our QLE assumptions, LD does not contribute to the approximated change in host or symbiont allele frequencies, and the approximation for change in LD reveals that it will remain negligibly small (derivation in Appendix A).

#### Sanctions with recognition

Finally, consider a model in which hosts have loci for symbiont recognition (with alleles *R* and *r*) and for sanctions (*H* and *h*), and symbionts have loci for signaling (*S* and *s*) and cooperation (*M* and *m*). Hosts initiate symbiosis only with symbionts carrying a signaling allele compatible with the hosts’ genotype at the recognition locus (i.e., *S* with *R* or *s* with *r*), as in the partner recognition model. However, hosts are also able to sanction if they carry the *H* allele at the sanctioning locus, as in the host sanctions model.

As in the host recognition model, to develop a tractable model we assume that the costs and benefits of interaction are small, that there is no epistasis, and that there is free recombination between loci. The payout values for each possible combination of host and symbiont genotypes (Table 3) then lead to the following approximations of change in the allele frequency at each locus in each species. For the host, these are

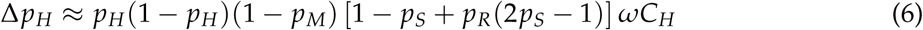

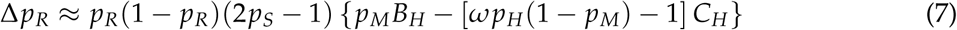

and for the symbiont

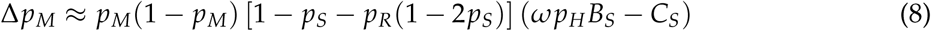

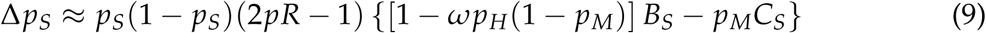

**Table 3:** Host and symbiont payouts under the model of recognition with sanctions.

| Symbiont | Host |  |  |  |
| --- | --- | --- | --- | --- |
|  | HR | Hr | hR | hr |
| *Host payout* |  |  |  |  |
| MS | $B_H - C_H$ | 0 | $B_H - C_H$ | 0 |
| Ms | 0 | $B_H - C_H$ | 0 | $B_H - C_H$ |
| mS | $-(1 - \omega)C_H$ | 0 | $-C_H$ | 0 |
| ms | 0 | $-(1 - \omega)C_H$ | 0 | $-C_H$ |
| *Symbiont payout* |  |  |  |  |
| MS | $B_S - C_S$ | 0 | $B_S - C_S$ | 0 |
| Ms | 0 | $B_S - C_S$ | 0 | $B_S - C_S$ |
| mS | $(1 - \omega)B_S$ | 0 | $B_S$ | 0 |
| ms | 0 | $(1 - \omega)B_S$ | 0 | $B_S$ |

As in the partner recognition model, the approximations for change in allele frequencies do not include terms for LD between host loci or between symbiont loci, meaning that LD does not affect the approximated change in allele frequencies for either species — and the approximations for change in LD in both species indicate that LD will remain negligible (see Appendix).

### Individual-based simulations

The approximations made to derive the analytic results may limit these models’ generality, and evaluation of equilibrium conditions provides a limited perspective given that few real biological communities are at an evolutionary equilibrium (Thompson 2013). Moreover, modeling dynamics in a single panmictic population misses the potential for divergence among geographically structured populations, which can be an important mechanism for diversification in coevolutionary systems (Nuismer et al. 1999; Thompson 2005; Yoder and Nuismer 2010; Thompson 2013). To account for a broader range of parameter space such as stronger fitness effects of mutualism, to evaluate results at non-equilibrium conditions, and to examine the effects of geographic population structure and diversification, we constructed an individual-based simulation of coevolution between hosts and symbionts in a metapopulation of sites linked by migration. (Parameters are given in Table 4; simulation scripts have been deposited in the Dryad Digital Repository, http://dx.doi.org/10.5061/dryad.p2s02; Yoder and Tiffin 2017)

**Table 4:** Parameter values for the individual-based simulations.

| Parameter<sup>1</sup> | Host | Symbiont |
| --- | --- | --- |
| *For metapopulation structure* |  |  |
| $K$, per-site population size | $U(20, 200)$ | $U(200, 2000)$ |
| $m$, among-site migration rate | $U(0, 0.05)$ | $U(0, 0.05)$ |
| *For interaction payouts* |  |  |
| $C$, cost of symbiosis | $U(0.01, 0.1)$ | $U(0.001, 0.1)$ |
| $B$, benefit of symbiosis | $C_H + U(0.01, 0.1)$ | $C_S + U(0.01, 1)$ |
| $\omega$, effectiveness of sanctions | $U(0.01, 1)$ | $U(0.01, 1)$ |
| *For genetics* |  |  |
| $r$, recombination rate | $U(0, 0.5)$ | $U(0, 0.5)$ |
| $\mu$, mutation rate | $10^{-6}$ | $10^{-6}$ |

<sup>1</sup> Parameters are either point values, or drawn from a uniform distribution with range $U(min, max)$ .

The simulation follows the change in allele frequencies in 50 populations of *K*_*i*_ haploid individuals of each species *i*, linked by migration at a rate of *m*_*i*_. We chose parameters to ensure that the interaction would be commensal or mutualistic (all *B*_*i*_ ≥ *C*_*i*_); and that symbionts would usually have larger population sizes than hosts and experience greater benefits from symbiosis. These asymmetries are seen in many mutualistic symbioses. The simulation starts by randomly creating individuals’ genotypes of one or two loci (depending on the model simulated) based on starting allele frequencies drawn from an approximation of the allele frequency spectrum for a standard neutral coalescent model at equilibrium (Ganapathy and Uyenoyama 2009); note that this means the simulations do not address conditions under which a new allele can invade a population, but the expected fate of genetic variants that are already segregating in a population when coevolution begins. After creation of the starting populations, the simulation proceeds through a generational cycle of migration among populations, interaction between hosts and symbionts in each population, and finally mating within populations with mating success determined by outcome of the host-symbiont interactions.

#### Migration

A proportion *m*_*i*_ of the individuals in each population are selected at random to join a global migrant pool, which are then distributed at random back among the populations.

#### Coevolutionary selection

Within each population, hosts interact with randomly-drawn symbionts. All hosts interact with symbionts, while symbionts that are not drawn for interaction are lost from the population. Each individual’s payout from the interaction is then determined by its genotype and the genotype of its host or symbiont, following one of the models outlined above. Finally, fitness is calculated for each individual as the payout of the interaction plus a value drawn at random from a normal distribution with mean = 1 and standard deviation = 0.1. These fitness values are then used to determine the probability of reproduction in the next step.

#### Mating

Mating occurs between pairs of hermaphroditic individuals of each species. Pairs of individuals are drawn at random from the same population, with replacement, and with the probability of being drawn scaled by each individual’s fitness value from the prior step. Each mating produces one offspring, with genotypes at each locus drawn from the parental genotypes. In two-locus species, recombination between loci occurs with probability *r*_*i*_, and mutation from one allele to the alternate allele occurs with probability *μ*_*i*_ for each locus. Mating continues until *K*_*i*_ offspring are created, at which time the offspring replace their parents to begin the next generation.

We ran 500 simulations for each of the three models as well as 500 simulations in which hosts and symbionts do not interact, which provide an expectation for evolution in the absence of the mutualism. Unless stated otherwise, we summarized simulation results after 1,000 generations, well past the time at which among-site variation in allele frequencies stabilized.

## Results

### Analytic models

We solved for equilibria in each of the three analytic models (sanctions only, recognition only, and sanctions with recognition) to identify conditions that maintain variation in host or symbiont loci, and that maintain mutualism (i.e., the frequency of symbiont cooperation, *p*_*M*_ *>* 0. Full details of these analyses are given in Mathematica notebooks, available in the Dryad Digital Repository, http://dx.doi.org/10.5061/dryad.p2s02 (Yoder and Tiffin 2017).

#### Sanctions

The sanctions model has no stable equilibria that maintain variation in either sanctions or cooperation (Figure 1). There are locally unstable equilibria when *p*_*M*_ = 0 and *p*_*H*_ is equal to either 1 or 0; and when *p*_*M*_ = 1 at any value of *p*_*H*_. Although variation is not maintained at either sanctions or cooperation loci, the rates of change in the frequencies of sanctioning and cooperation alleles are very low whenever *p*_*M*_ or *p*_*H*_ are near 1, meaning that it may take considerable time for these alleles to become fixed.

**Figure 1:**
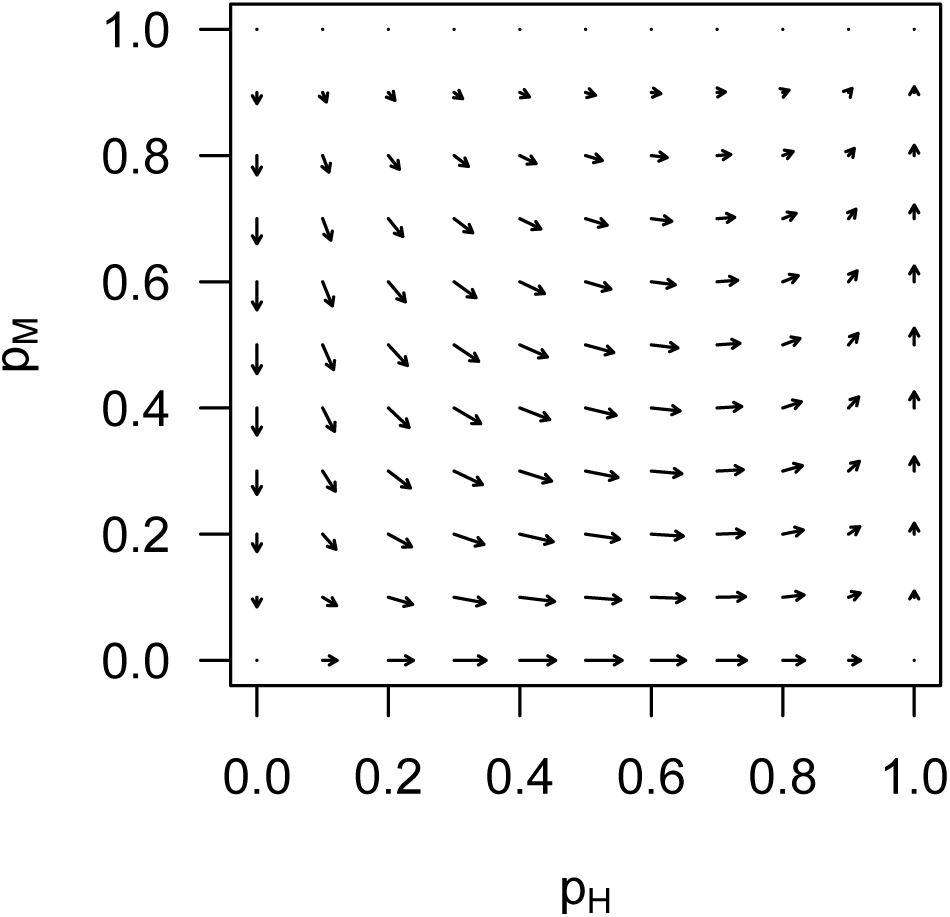
Dynamics of the host sanctions model. Vector-field plot indicating magnitude and direction of change in the frequency of host sanctions (*p*_*H*_) and symbiont cooperation (*p*_*M*_) alleles at given starting frequencies, with *C*_*H*_ = *C*_*S*_ = 0.25, *B*_*H*_ = *B*_*S*_ = 0.5, and *ω* = 0.75.

#### Partner recognition

In the partner recognition model there are locally unstable equilibria that maintain variation, with cyclical dynamics, at recognition and signaling loci (*pR* = *pS* = 0.5), but only when the symbiont cooperation allele (*M*) is fixed or lost (Figure 2). There are also unstable equilibria that maintain variation in cooperation (i.e., *M* is at intermediate frequency), but only when the host’s recognition locus and symbiont’s signaling locus are fixed for incompatible alleles. In other words, the system can maintain variation in recognition, but only when that recognition has no consequences for fitness; and it can maintain variation for cooperation, but only when symbiosis is never initiated. Under these conditions variation maintained at mutualism-related loci is effectively neutral, and would be lost via drift in finite populations.

**Figure 2:**
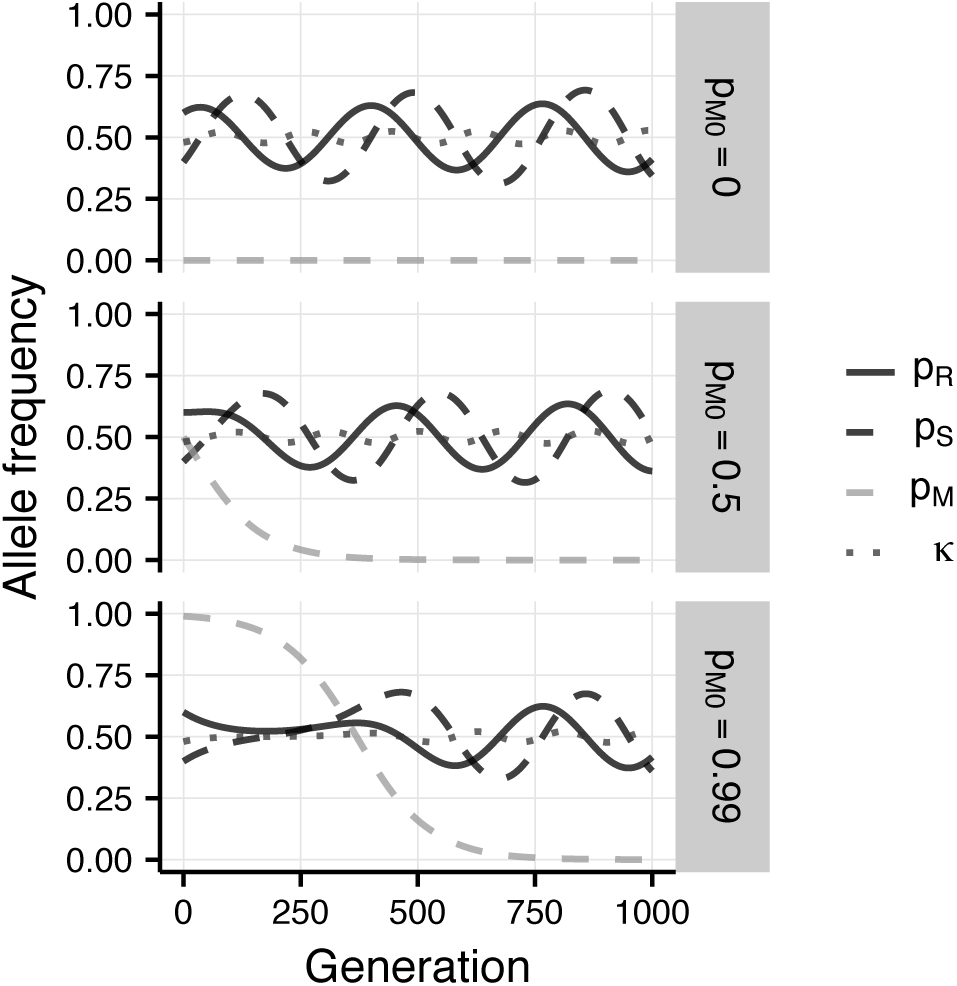
Dynamics of partner recognition. Allele frequencies over time at host recognition (*p*_*R*_, solid dark lines) and symbiont signaling (*p*_*S*_, dashed dark lines) loci, allele frequency at the symbiont cooperation locus (*p*_*M*_, dashed gray lines), and host-symbiont compatibility, (*κ*, dotted lines), with intial frequency of the symbiont cooperation allele *p*_*M*0_ = 0, *p*_*M*0_ = 0.5, or *p*_*M*0_ = 0.99, and with *C*_*H*_ = *C*_*S*_ = 0.025 and *B*_*H*_ = *B*_*S*_ = 0.05.

Variation at signaling loci is maintained by inverse frequency dependent selection when the cooperation allele is lost from the population (*p*_*M*_ = 0). However, this is not surprising given that when *M* = 0 the system is effectively a host-parasite system, which have been repeatedly shown to maintain variation with a 2-allele system determining host recognition (Dieckmann et al. 1995; Agrawal and Lively 2002). We can examine these dynamics in terms of host-symbiont *compatibility*, or the probability that a randomly-drawn host and symbiont will carry compatible recognition and signaling alleles, defined as *κ* = *p*_*R*_ *p*_*S*_ + (1 - *p*_*R*_)(1 - *p*_*S*_). This probability remains close to 0.5 even as recognition and signaling allele frequencies cycle (Figure 2).

Perhaps the most important feature of the model of partner recognition model is that change in the frequency of the symbiont cooperation allele, Δ*p_M_*, is ≤ 0 for all reasonable parameter values (Equation 4; see also the “recognition” Mathematica notebook at http://dx.doi.org/10.5061/dryad.p2s02; Yoder and Tiffin 2017). That is, there is no condition under which a cooperative symbiont allele (*M*) will increase in frequency. This means that partner recognition alone cannot select for greater frequency of cooperation, from any starting frequency (Figure 2).

#### Sanctions with recognition

The model of partner recognition with host sanctions has equilibria that maintain variation at both host recognition and symbiont signaling loci. Still, the only locally stable equilibria at which hosts or symbionts maintain variation in sanctioning or cooperation also have hosts and symbionts fixed for incompatible recognition/signaling alleles — conditions ensuring that symbiosis is never initiated. Unstable equilibria exist where there is variation in host sanctions or symbiont cooperation without complete loss of host-symbiont compatibility, and where cyclical dynamics can occur at the host recognition and symbiont signaling loci (Figure 3, central panel). These depend on starting allele frequencies, the cost and benefit of mutualism to each species, and particularly the effectiveness of sanctions.

**Figure 3:**
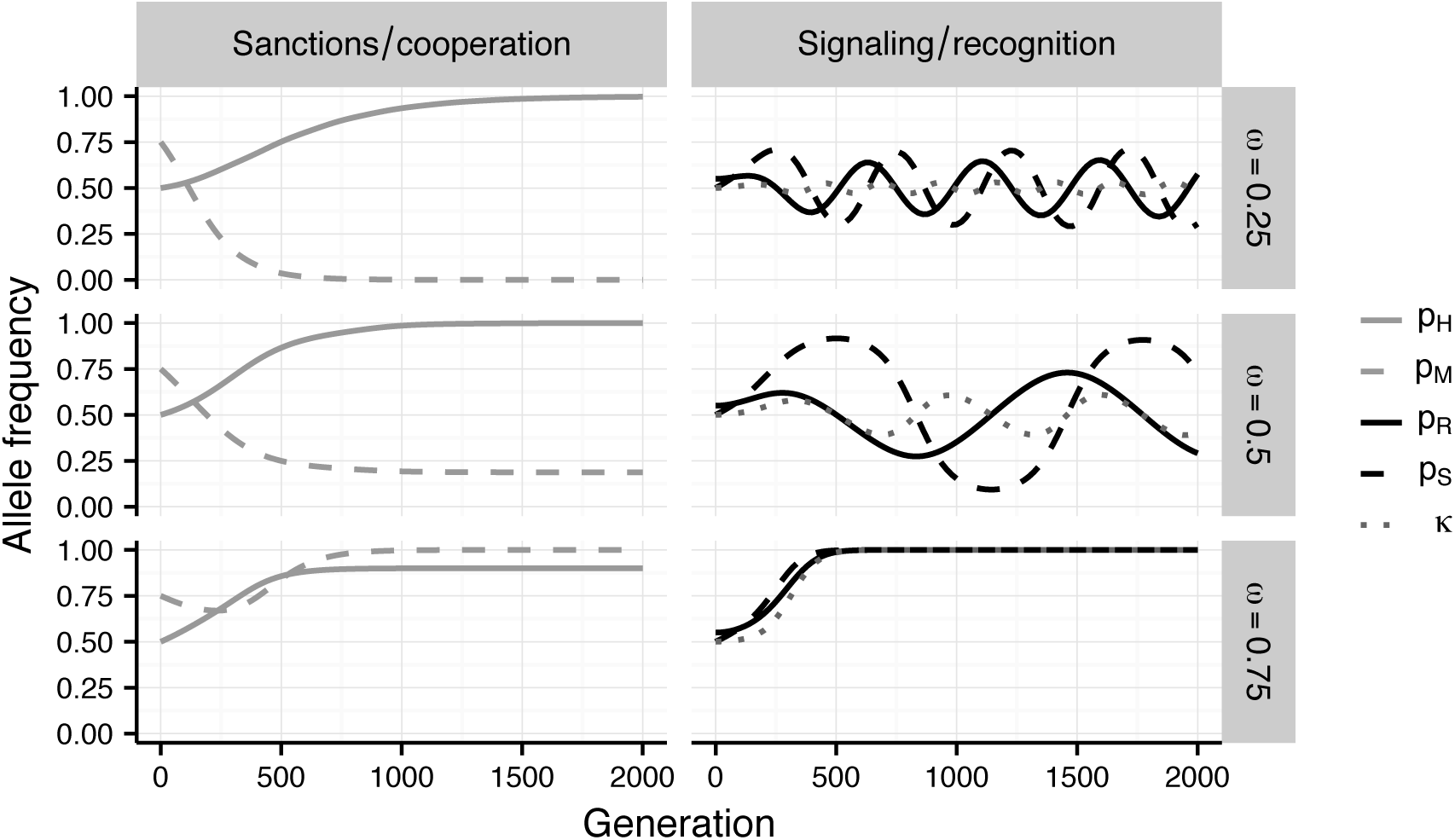
Dynamics of host sanctions with partner recognition. Frequency of the host sanctions allele (*p*_*H*_, left panels, solid lines) and symbiont cooperation allele (*p*_*M*_, left panels, dashed lines) or frequency of the host recognition allele (*p*_*R*_, right panels, solid lines) and symbiont signaling allele (*p*_*R*_, right panels, dashed lines) with host-symbiont compatibility (*κ*, right panels, dotted lines), when the effectivness of sanctions *ω* = 0.25 (top), 0.5 (middle), or 0.75 (bottom). For all scenarios, the initial frequency of the host sanctions allele *p*_*H*_ = 0.5, host recognition allele *p*_*R*_ = 0.55, symbiont cooperation *p*_*M*_ = 0.75, symbiont signaling *p*_*S*_ = 0.5, *C*_*H*_ = *C*_*S*_ = 0.025 and *B*_*H*_ = *B*_*S*_ = 0.05.

If sanctions are not very effective (i.e., *ω* is low) then the symbiont cooperation allele *M* may be lost even as the host sanctions allele *H* become fixed; loss of cooperative symbionts leads to inverse frequency-dependent cycling at the host recognition and symbiont signaling loci, as seen in a host-pathogen model (Figure 3, upper panels). With more effective sanctions, symbiont cooperation can be maintained at low frequency, while inverse frequency-dependent cycles at the signaling and recognition loci still occur — reflecting the fact the symbiosis is, on average, detrimental to host fitness so long as fewer than half of symbionts are cooperative (Figure 3, middle panels).

When sanctions are sufficiently effective, variation can be maintained at the host sanctions locus once symbiont cooperation is fixed and the recognition/signaling loci become fixed for compatible alleles (i.e., *R* and *S*; Figure 3, lower panels). This parallels results from the model of host sanctions alone (Figure 1), in which fixation of symbiont cooperation reduces Δ*p_H_* to zero (Equation 1). Indeed, when the signaling and recognition loci are fixed for compatible alleles (*κ* = 1), the model of sanctions with recognition behaves like the model of sanctions alone.

Meanwhile, the relative fitness of each symbiont signaling allele depends not only on the frequency of host recognition alleles (*p*_*R*_), but also on the frequency of symbiont cooperation (*p*_*M*_), the frequency of sanctioning hosts (*p*_*H*_), and the degree to which sanctions reduce the fitness of non-cooperating symbionts (*ω*). This is apparent from the approximate expression for change in the frequency of the symbiont signaling allele, Δ*p_S_* which has an unstable equilibrium when *p*_*H*_ = 1 and *C*_*S*_/*B*_*S*_ = 1 - *ω*(1 - *p*_*M*_) (Equation 9).

Host-symbiont compatibility, *κ*, increases most rapidly when most symbionts are cooperative, when sanctioning hosts are more common, and when hosts have near-maximum variation at the recognition locus (*p*_*R*_ is near 0.5) but symbionts are nearly fixed for one signaling allele (Figure 4, dark-shaded regions). It decreases most rapidly when hosts are mostly unable to sanction, cooperative symbionts are at lower frequency (*p*_*M*_ *<* 0.5), and hosts mostly carry a recognition allele compatible with the more-common symbiont allele (Figure 4, light-shaded regions).

**Figure 4:**
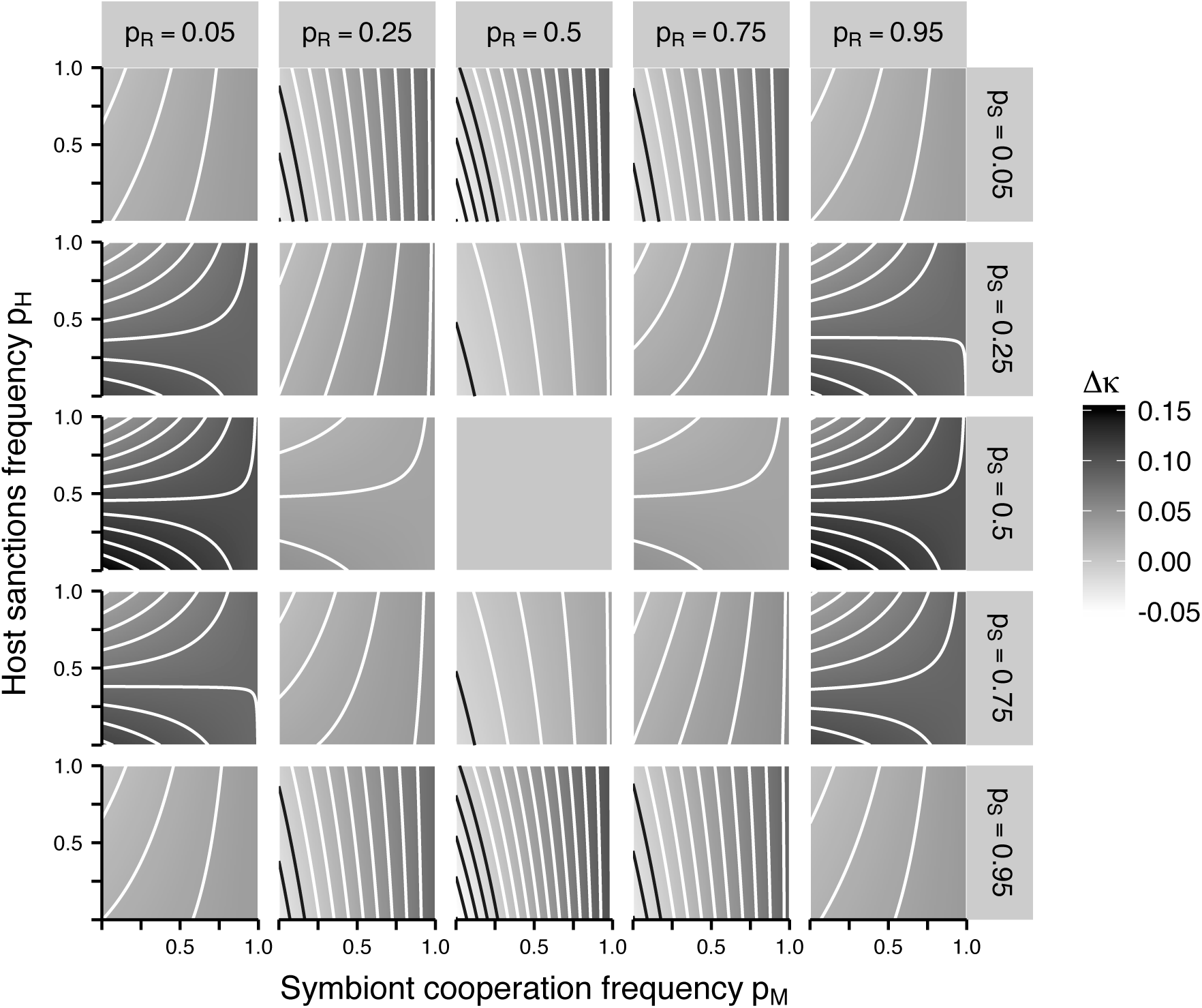
Analytical results showing per-generation change in host-symbiont compatibility Δ*κ*, for different starting frequencies of the symbiont cooperation allele, *p*_*M*_, and host sanctions allele, *p*_*H*_; and for different starting frequencies of the host recognition allele, *p*_*R*_, and symbiont signaling allele, *p*_*S*_. Darker shading indicates greater values of Δ*κ*; contour lines are at intervals of 0.01, with white lines indicating values of Δ*κ >* 0. For all panels *ω* = 0.75, *B*_*S*_ = *B*_*H*_ = 0.75, and *C*_*S*_ = *C*_*H*_ = 0.25.

### Individual-based simulations

#### Sanctions

After 1,000 generations, individual-based simulations of the sanctions model ended with the host sanctioning allele at significantly higher frequency than expected from neutral simulations (Figure 5A; *p <* 1 × 10^−6^, t-test on arcsine-transformed values). This is consistent with the predictions of the analytic model (Figure 1). Still, the sanctioning allele, *H*, became fixed in only 21% of simulations, and there was considerable overlap in the range of frequencies seen for the sanctioning allele and that for neutral alleles. Sanctioning hosts were more common when costs of symbiosis for the hosts were greater (Spearman’s *ρ*_*p_H_*, *C*_*H*__ = 0.25, *p <* 1 × 10^−6^) and when the benefits of symbiosis were greater (*ρ*_*p_H_*, *B*_*H*__ = 0.15, *p <* 0.001).

**Figure 5:**
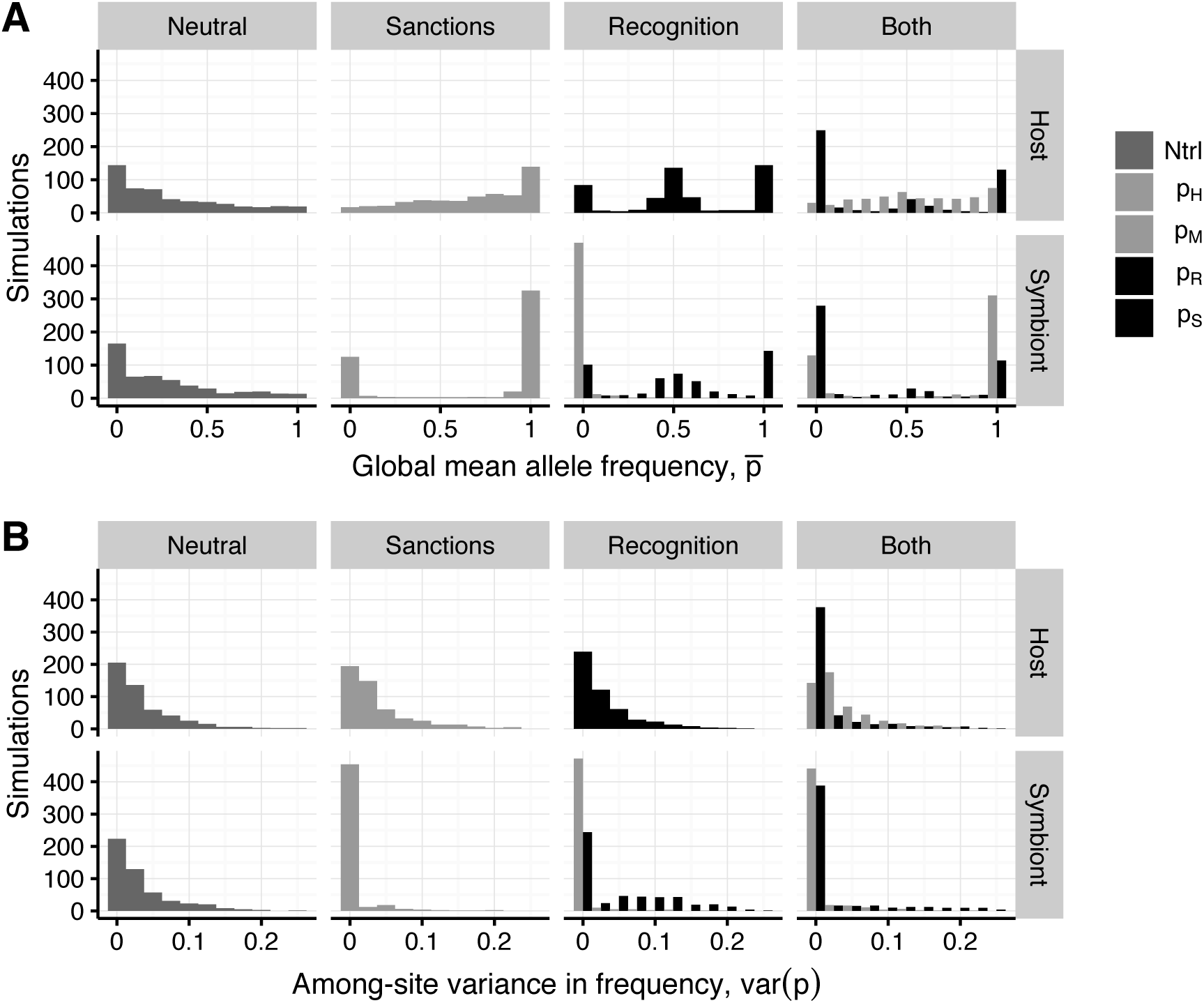
(A) Mean within-population allele frequencies and (B) among-site variation in local allele frequency after 1,000 generations of coevolution with no interaction (neutral), or with each of the three models of host-symbiont interaction. Results are from 500 replicate simulations with parameters given in Table 4.

Sanctions resulted in higher symbiont cooperation (Figure 5A), again consistent with the predictions of the analytic model (Figure 1). The cooperation allele, *M*, rose to significantly higher frequency than neutral alleles (t-test *p <* 1 × 10^−6^), and became fixed in 59% of sanctions simulations. However, cooperation was lost in 21% of simulations. Cooperative symbionts were less common when the cost of symbiosis was higher (*ρ*_*p_M_*, *C*_*S*__ = *-*0.20, *p <* 1 *×* 10^−5^); in contrast, cooperative symbionts were more common when the benefits were higher (*ρ*_*p_M_*, *B*_*S*__ = 0.23, *p <* 1 *×* 10^−6^), and when host sanctions were more effective (*ρ*_*p_M_*, *ω*_ = 0.53, *p <* 1 *×* 10^−6^).

To examine geographic differentiation among sites, we calculated the among-site variance in allele frequencies for each replicate simulation (Figure 5B). In simulations of the sanctions model, among-site variance in the frequency of the host sanctions allele was very similar to that seen for neutral simulations. The symbiont cooperation locus, however, had much lower among-site variation (*var*(*p*_*M*_) = 0 in 80% of simulations) than neutral loci (*var*(*p*) = 0 in 20% of neutral simulations), reflecting the high proportion of simulations in which symbiont cooperation became fixed (Figure 5A).

#### Partner recognition

In simulations of partner recognition alone, symbiont cooperation was lost in the vast majority of simulations (79%; Figure 5A), as predicted by the analytic model (Figure 2). Consistent with the cyclical dynamics predicted by the analytic model, host recognition alleles were more likely to be at intermediate frequencies than neutral alleles (0.4 *< p*_*R*_ *<* 0.6 in 42% of simulations, versus 12% for neutral alleles), and the same was true for symbiont signaling alleles (0.4 *< p*_*S*_ *<* 0.6 in 28%, versus 10% for neutral alleles). Signaling and recognition alleles were also often fixed or lost, but could resume cyclical dynamics when variation was reintroduced by mutation (Figure 6).

**Figure 6:**
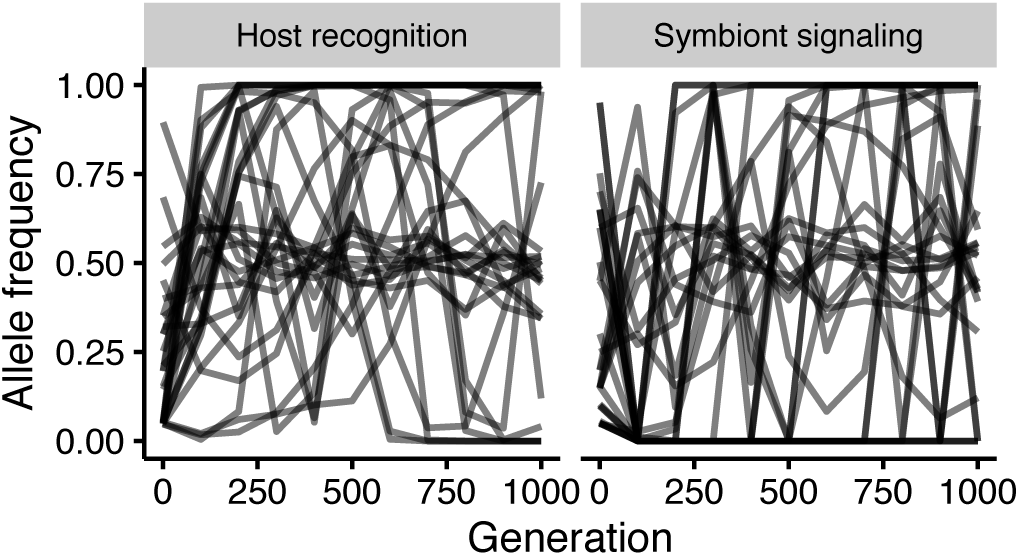
Cyclical dynamics in individual-based simulations of recognition. Plots of allele frequency over time at the host recognition locus (*p*_*R*_; left) and symbiont signaling locus (*p*_*S*_; right), in a sample of 30 replicate simulations.

Patterns of among-site variation at the symbiont signaling and host recognition loci were broadly similar to those seen for neutral loci (Figure 5B). However, because the symbiont cooperation allele was so often lost, among-site variation in the frequency of the the symbiont cooperation allele was much lower than either seen for either the symbiont signaling alleles or neutral alleles (*var*(*p*_*M*_) = 0 in 79% of simulations, *var*(*p*_*S*_) = 0 in 42%; *var*(*p*) = 0 in 22%).

#### Sanctions with recognition

Simulations of sanctions with recognition resulted in, on average, somewhat lower frequency of sanctioning hosts than simulations of sanctions alone (t-test on arcsin-transformed data, *p <* 1 × 10^−6^), and were less likely to have sanctions fixed (*p*_*H*_ = 1 in 11% of simulations of sanctions with recognition, compared to 21% with sanctions alone). As in simulations of sanctions alone, sanctioning hosts were more common when the cost of symbiosis was higher (*ρ*_*p_H_*, *C*_*H*__ = 0.36, *p <* 1 *×* 10^−6^) and when the benefits were greater (*ρ*_*p_H_*, *B*_*H*__ = 0.21, *p <* 1 *×* 10^−5^).

The fate of symbiont cooperation in simulations of sanctions with recognition was also similar to results from the simulations of sanctions alone. In 56% of simulations the cooperation allele went to fixation, though it was lost in 21%. Also as in the simulations of sanctions alone, cooperative symbionts were more common when costs of symbiosis were lower (*ρ*_*p_M_*, *C*_*S*__ = *-*0.22, *p <* 1 *×* 10^−6^), when benefits were greater (*ρ*_*p_M_*, *B*_*S*__ = 0.20, *p <* 1 *×* 10^−5^), and when host sanctions were more effective (*ρ*_*p_M_*, *ω*_ = 0.61, *p <* 1 *×* 10^−6^).

The frequency of host recognition alleles was strongly and positively correlated with the frequency of symbiont signaling alleles at all timepoints (at generation 1,000, *ρ* = 0.80 for *p*_*R*_ and *p*_*S*_, *p <* 1 × 10^−6^), and recognition and signaling alleles were at intermediate frequency in fewer simulations of sanctions with recognition (0.4 *< p*_*R*_ *<* 0.6 in 12% of simulations; 0.4 *< p*_*S*_ *<* 0.6 in just 9%) compared to simulations of recognition alone. This is consistent with hosts and symbionts converging on compatible signaling and recognition alleles, and maintaining them at high frequency — something also seen in the analytical model when sanctions were effective. Indeed, in simulations of sanctions with recognition, host-symbiont compatibility (*κ*) was generally much higher than in the model of recognition alone (Figure 7, *p <* 1 × 10^−6^ in a t-test on arcsin-transformed values).

**Figure 7:**
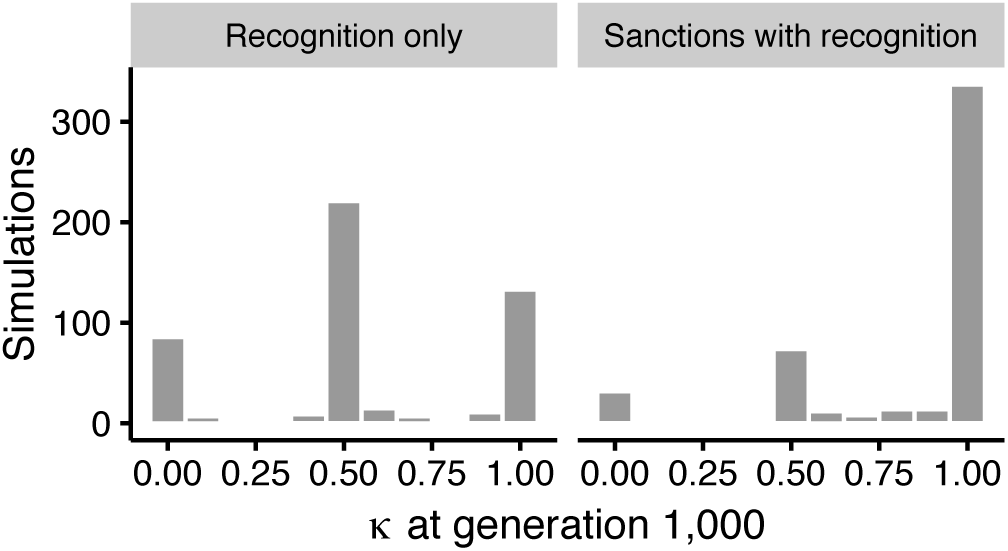
Host-symbiont compatibilty, *κ*, after 1,000 generations of coevolution in simulations of recognition alone (left) or sanctions with recognition (right).

In simulations of sanctions with recognition the two mechanisms of interaction (i.e., sanctions/cooperation and recognition/signaling) evolved in response to each other as predicted by the analytic model (Figures 3 and 4). Hosts and symbionts were more compatible when there were more sanctioning hosts at early time points in the simulation, but this correlation decreased over time, possibly due to fixation of the sanctioning allele in many replicates (tests on arcsine-transformed values; at generation 100, Spearman’s *ρ*_*p_H_*, *κ*_ = 0.25, *p <* 1 × 10^−6^; at generation 500, *ρ*_*p_H_*, *κ*_ = 0.17, *p* = 0.001; at generation 1,000, *ρ*_*p_H_*, *κ*_ = 0.05, *p* = 0.30).

Notably, simulations of sanctions with recognition maintained significantly more among-site variation at the sanctions locus than simulations of sanctions alone (t-test on arcsin-transformed values; *p* = 0.005), and maintained much less among-site variation at the host recognition locus than simulations of recognition alone (t-test *p <* 1 × 10^−6^). Simulations of sanctions alone and sanctions with recognition both resulted in very low among-site variation at the symbiont cooperation locus, with *var*(*p*_*M*_) = 0 in 76% and 70% of simulations, respectively.

## Discussion

Variation in the quality of mutualistic partners is widely observed in natural systems, (Pellmyr and Huth 1994; Herre and West 1997; Holland et al. 1999; Simms and Taylor 2002; Ness et al. 2006; Heath and Tiffin 2009; Hoeksema 2010; reviewed by Heath and Stinchcombe 2014), in spite of the fact that mechanisms that stabilize mutualisms against cheating should remove such variation over time (Axelrod and Hamilton 1981; Doebeli and Knowlton 1998; West et al. 2002b; Kopp and Gavrilets 2006; Yoder and Nuismer 2010). Genetic variation can be maintained in the face of selection by mutation-selection balance (Foster et al. 2006) or by drift and migration among spatially structured populations (Thompson et al. 2013; Heath and Stinchcombe 2014), but none of the forms of coevolutionary selection typically expected between mutualists are expected to maintain variation.

Our model of symbiotic mutualism in which hosts recognize symbiont signals and sanction non-cooperative symbionts shows how coevolutionary selection between mutualists can both stabilize mutualism and maintain variation in its outcomes. Neither sanctions nor partner recognition alone maintain mutualism and variation in mutualism outcomes. However, in a model that includes both mechanisms, variation can be maintained either because hosts and symbionts vary in sanctioning ability and cooperation, or because they vary in their signaling/recognition compatibility (Figure 3). Our individual-based simulations corroborate the prediction from the analytic model that the sanctions and recognition systems interact by altering the coevolutionary conditions each genetic system faces (Figures 5, 7).

### Sanctions versus recognition, solo and in concert

Similar to previous models (Trivers 1971; Axelrod and Hamilton 1981; Bull and Rice 1991; West et al. 2002a; West et al. 2002b; Foster and Wenseleers 2006), ours finds that host sanctions maintain cooperative symbionts at high frequency (Figures 1, 5). The advantage of non-cooperation may cause the frequency of cooperative symbionts to decrease, but lower frequency of cooperation increases the relative fitness of sanctioning hosts — and once sanctions are sufficiently common, the frequency of cooperative symbionts increases to fixation (Figure 1).

On the other hand, when partner signals and cooperation are determined by unlinked loci, partner recognition alone is unable to select for greater frequency of cooperative symbionts (Equation 4), and when cooperative symbionts are lost selection favors hosts compatible with whichever symbiont signaling allele is less common (Figure 2). Because hosts gain no benefit from the symbiosis, this situation is effectively the loss of mutualism, and it creates coevolutionary cycles in the frequency of signaling and recognition alleles (Figures 2, 6), similar to what is seen in host-pathogen systems (Dieckmann et al. 1995; Agrawal and Lively 2002; M’Gonigle and Otto 2011).

In contrast to these simpler systems, when hosts both selectively initiate symbiosis based on recognition of symbiont signals and sanction non-cooperative symbionts, the system can maintain variation in at host sanctions and recognition loci, and at symbiont cooperation and signaling loci (Figure 3, middle panels). However, this occurs only when sanctions have intermediate effectiveness; if sanctions are less effective, the same starting conditions and interaction payout result in loss of cooperation (Figure 3, top panels); whereas if sanctions are stronger, symbiont cooperation becomes fixed and recognition and signaling loci fix for compatible alleles (Figure 3, bottom panels). Once cooperation is fixed, the host sanctioning allele is effectively neutral, and variation at that locus is expected to be lost to drift. If there were a cost to simply maintaining the ability to sanction, we might expect that sanctioning hosts would become less common again, creating an opportunity for non-cooperative symbionts to re-emerge via mutation — creating cyclical dynamics over longer time periods than we consider here. Because fixation of symbiont cooperation eliminates the source of selection favoring sanctioning, it has been proposed that re-introduction of non-cooperative partners, or interaction with multiple partner species that vary in their cooperativeness, is necessary to maintain sanctions in mutualist populations over the long term (Foster et al. (2006)).

Our simulations of sanctions with recognition show somewhat less frequent fixation of the host sanctioning allele and more among-site variation in its frequency than simulations of sanctions alone (Figure 5A, 5B). These outcomes are connected to the fact that when hosts can sanction non-cooperative symbionts there is less selective advantage to avoiding symbiosis, and hosts and symbionts converge on compatible recognition and signaling alleles (Figure 7). These results recall those from models of mutualism based on economic contract theory, which propose that sanctioning is often best understood not as a specific adapation to minimize the cost of interaction with non-cooperative partners (Weyl et al. 2010; Archetti et al. 2011a; Archetti et al. 2011b), but as pre-existing characteristics of the mutualists that provide partner fidelity feedback by positively responding to cooperative symbionts (Archetti et al. 2011a; Frederickson 2013). Under this thinking, floral abortion in response to pollinator overexploitation in brood pollination mutualisms is a repurposing of plants’ response to floral damage, and legumes’ reduced allocation to underproductive root nodules may be due to adaptations for root growth in soil with heterogenous nutrient content (Pellmyr and Huth 1994; Kiers et al. 2006; Weyl et al. 2010; Batstone et al. 2017). In our model, interaction with symbionts of varying quality leads to higher frequency, and often fixation, of sanctioning hosts (Figures 3, 5A). High frequency of sanctioning hosts then relaxes selection for host recognition alleles that prevent symbiosis, resulting in higher host-symbiont compatibility (Figures 4, 7). This recapitulates the result of Archetti *et al.*, (2011a) that hosts offering the right “terms” to symbionts need not screen for cooperative indivdiuals prior to initiating the interaction. (In our model, the “terms” offered to symbionts would be that hosts will not sanction if symbionts cooperate.)

Although few well-studied mutualisms involve two haploid partners, as in our models, we do not believe that relaxing this assumption would change our conclusions. M’Gonigle and Otto (2011) modeled the effect of host and symbiont ploidy in a matching-alleles model similar to the system we consider, and found that diploidy hosts were better able to recognize and resist both haploid and diploid parasites. This suggests that a diploid version of our partner recognition model would see hosts better able to evade symbiosis when cooperative symbionts are rare, but such evasion does not select for more cooperative symbionts (Figure 2). A diploid model of host sanctions could allow more continuous variation in sanctioning effectiveness and symbiont cooperation, but the fundamental dynamic of sanctions selecting for more cooperative symbionts should remain (Figure 1, 3).

### Cooperation and communication in mutualism

Our results suggest that multiple genetic mechanisms, which may experience very different forms of coevolutionary selection, may contribute to the evolution of cooperating species. Evaluating this prediction will require experiments that explicitly separate the exchange of benefits from the initiation of a mutualism (e.g., Regus et al. 2014; Althoff 2016; Powell and Doyle 2016). Population genetic studies that test for different signals of selection at loci with different roles in mutualism may also provide insight into the long-term selection that has acted on different stages of an interaction (e.g., Paape et al. 2013; Bonhomme et al. 2015; Yoder 2016).

There is evidence in many mutualisms for communication between partners that is based on traits separate from the rewards provided to (or withheld from) those mutualists (Svensson et al. 2005; Edwards et al. 2006b; Okamoto et al. 2007; Soler et al. 2011; Svensson et al. 2016). The system in which the relationship between partner signals and response to partner performance is best understood may be the symbiotic mutualism of legumes and nitrogen-fixing rhizobial bacteria. Legumes sanction ineffective rhizobia (Kiers et al. 2003; Kiers et al. 2006; Regus et al. 2014) or preferentially allocate resources to nodules hosting effective rhizobia (Heath and Tiffin 2009; Batstone et al. 2017), but they also respond to molecular signals from rhizobia as they establish symbiosis (Triplett and Sadowsky 1992; Oldroyd et al. 2011). At the level of quantitative phenotypes, host-rhizobium compatibility is at least partly independent of variation in mutualism outcomes (Triplett and Sadowsky 1992; Bena2005a; Heath and Tiffin 2009; Grillo et al. 2016; Powell and Doyle 2016).

Members of legume gene families associated with pathogen recognition are also implicated in legume-rhizobium compatibility (Yang et al. 2010; Young et al. 2011), and some legume genes with roles in the symbiosis show elevated nucleotide diversity and geographic differentiation consistent with frequency-dependent selection (Yoder 2016). By contrast, rhizobial genes producing nodule initiation factors and type III effector genes, both of which can be recognized by hosts, have reduced diversity (Bailly et al. 2006; Kimbrel et al. 2013). Another complicating factor is that many rhizobia species have genes involved in signaling and nitrogen fixation physically linked in a “symbiosis island” (e.g., Sullivan and Ronson 1998; Laguerre et al. 2001; Parker 2012), which might reduce the opportunity for separate evolution of signaling and cooperation. Still, there are examples of rhizobial genes mediating host recognition that exhibit signs of negative frequency-dependent selection when they are not in close linkage with nitrogen fixation genes (Bailly et al. 2006), and genes involved in both signaling and nitrogen fixation show signs of elevated horizontal gene transfer (Bailly et al. 2006; Sun et al. 2006; Epstein et al. 2012; Parker 2012). Sequence conservation at rhizobial signaling genes may also be explained by one of our key results, in which hosts’ ability to sanction allows hosts and symbionts to converge on compatible alleles at recognition and signaling loci (Figures 3, 6) — such convergence should create selection to maintain compatibility.

## Conclusions

Mutualistic interactions require communication between potential partners as well as cooperation once the interaction is underway. Previous theory of mutualism has not, however, explicitly included both of these systems of interaction. Our model of a symbiotic mutualism incorporating host recognition of symbiont signals alongside host sanctions against non-cooperative symbionts proves to better reflect the apparent contradictions of empirical systems, maintaining mutualism and variation in interaction outcomes under conditions where neither system, on its own, can do so.

## Data archiving

Full derivation and analysis of our analytic models, and scripts for individual-based simulations, are in the Dryad Digital Repository at http://dx.doi.org/10.5061/dryad.p2s02.

## Acknowledgements

This work began as a project for the UMN Theory Under Construction discussion group. We also thank the Whitlock lab, attendees of the UBC Biodiversity Lunchtime Informal Seminar Series, Scott L. Nuismer, and two anonymous reviewers. This work was funded by a National Science Foundation award to PT (grant 1237993, N.D. Young, P.I.), and we developed and performed our simulations on computing systems maintained by the Minnesota Supercomputing Institute and Compute Canada’s Westgrid network. JBY is currently a postdoctoral fellow with the CoAdapTree Project, supported by Genome Canada with co-funding from Genome British Columbia and Genome Quebec (co-Project Leaders S.N. Aitken, S. Yeaman, and R. Hamelin).

### Appendix: Evolution of linkage disequilibrium

#### Change in LD in the partner recognition model

Under the QLE assumptions described in the main text and in the Mathematica notebooks in the Supplementary Information (github.com/jbyoder/mutualism-sanctions-recognition), we can approximate change in linkage disequilibrium (LD) between the symbiont’s mutualism and signaling loci, *δ*_*MS*_:

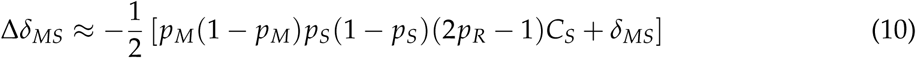

Δ*δ*_*MS*_ has the opposite sign of *δ*_*MS*_, and LD between the symbiont cooperation and signaling loci will evolve toward zero, unless -*p*_*M*_(1 - *p*_*M*_)*p*_*S*_(1 - *p*_*S*_)(2*p*_*R*_ - 1)*C*_*S*_ *< δ*_*MS*_ *<* 0, or 0 *< δ*_*MS*_ *<* -*p*_*M*_(1 - *p*_*M*_)*p*_*S*_(1 - *p*_*S*_)(2*p*_*R*_ - 1)*C*_*S*_. For parameter values meeting the assumptions of the approximation (small *C*_*S*_) this means that LD between symbiont loci will remain negligible.

#### Change in LD in the model of sanctions with recognition

The model of sanctions with recognition gives the following QLE approximations for change in LD between the host sanctions and recognition loci, *δ*_*HR*_

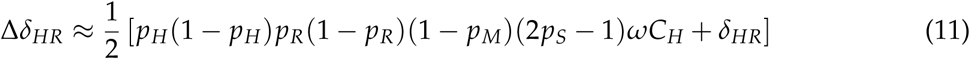

And for change in LD between the symbiont cooperation and signaling loci, *δ*_*MS*_

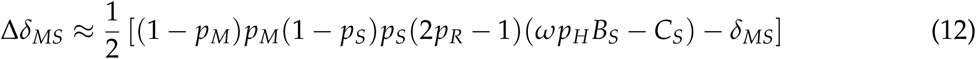

For the hosts, Δ*δ*_*HR*_ and *δ*_*HR*_ have opposite signs, and LD between the sanctions and recognition loci evolves toward zero, unless 0 *< δ*_*HR*_ *< p*_*H*_(1 - *p*_*H*_)*p*_*R*_(1 - *p*_*R*_)(1 - *p*_*M*_)(2*p*_*S*_ - 1)*ωC*_*H*_ or *p*_*H*_(1 - *p*_*H*_)*p*_*R*_(1 - *p*_*R*_)(1 - *p*_*M*_)(2*p*_*S*_ - 1)*ωC*_*H*_ *< δ*_*HR*_ *<* 0. In symbionts, LD between the cooperation and signaling loci evolves toward zero unless 0 *< δ*_*MS*_ *< p*_*M*_(1 - *p*_*M*_)*p*_*S*_(1 - *pS*)(2*p*_*R*_ - 1)(*ω p*_*H*_ *B*_*S*_ - *C*_*S*_) or *p*_*M*_(1 - *p*_*M*_)*p*_*S*_(1 - *pS*)(2*p*_*R*_ - 1)(*ω p*_*H*_ *B*_*S*_ - *C*_*S*_) *< δ*_*MS*_ *<* 0. In both species, the conditions required for the approximation (small cost, *C*_*i*_ and benefit, *B*_*i*_ terms for each species) make the values of LD in these ranges negligibly small.

